# Major antigenic differences in *Aeromonas salmonicida* isolates correlate with the emergence of a new strain causing furunculosis in Chilean salmon farms

**DOI:** 10.1101/2025.01.30.635334

**Authors:** Marcos Mancilla, Adriana Ojeda, Yassef Yuivar, Maritza Grandón, Horst Grothusen, Marcela Oyarzún, Alejandro Bisquertt, Juan A. Ugalde, Francisco Fuentes, Pablo Ibarra, Patricio Bustos

## Abstract

*Aeromonas salmonicida* is the etiological agent of furunculosis, a septicemic disease with high mortality rates affecting salmonids and other teleost species worldwide. Reviewing molecular diagnostic protocols for routine diagnostics, we realized that the amplification of the *vapA* target gene failed in some cases of furunculosis. Therefore, we hypothesized that the emergence of a new strain may be involved in recent outbreaks. In this work, we demonstrate that the *vapA* locus is absent in the new strain, which explains why it lacks the major membrane component VapA protein, a critical virulence factor. In addition, we found that the *vapA-*absent strain differs from its counterparts in outer membrane protein and lipopolysaccharide profiles, suggesting profound changes at the membrane structure level and in antigenic properties. These features along with sequence analysis information allowed us to infer that a complex genomic rearrangement, probably an indel encompassing the entire *vapA* locus, gave rise to this membrane phenotype. Although the causes for pathogen evolution and emergence were not fully elucidated, our results strongly suggest that the *vapA-*absent strain is responsible for a raising proportion of recent furunculosis cases, and that it may be related to a less virulent disease and a low serological response upon vaccination with the *A. salmonicida* antigen formulation currently used in Chile.

## 1. INTRODUCTION

*Aeromonas salmonicida* is a Gram-negative bacterium and etiological agent of furunculosis, a bacterial disease that affects reared fish, producing high economic impact in aquaculture worldwide (Bartkova et al., 2017b). Taxonomically, five subspecies of *A. salmonicida* have been described to be pathogenic for fish (Long et al., 2023). The presence of typical skin lesions referred to as furuncles, as well as ulcers, exophthalmia, hemorrhages, and septicemia are commonly seen in salmon infected by *A. salmonicida* subsp. *salmonicida* and result in acute mortality. Atypical subspecies, *i.e.*, *A. salmonicida* subsp. *achromogenes*, *A. salmonicida* subsp. *masoucida*, *A. salmonicida* subsp. *smithia*, and *A. salmonicida* subsp. *pectinolytica* were reported on the basis of biochemical differences such as reduced or slow pigmentation and growth at temperatures over 20 °C. They produce similar clinical signs but affect a broader number of hosts including carp (*Cyprinus carpio*), goldfish (*Carassius auratus*), and flounder (*Platichthys flesus*) (Austin et al., 1998; Wiklund and Dalsgaard, 1998), with the exception of *A. salmonicida* subsp. *pectinolytica*, which was originally isolated from polluted water (Pavan et al., 2000). In genomic terms, atypical *A. salmonicida* strains are less complex, harboring a single, large plasmid, but have enhanced plasticity capacity that is ascribed to a high number of insertion sequences (Nilsson et al., 2006; Vasquez et al., 2022). Although the genomes of atypical strains present *loci* coding genes for virulence factors such as flagella and type three secretion systems, it seems that pseudogenization has inactivated certain components, which could compromise the functionality of these virulence factors and eventually contribute to a more chronic course of the disease (Vasquez et al., 2022). Insofar as outbreaks in Chile were characterized in this respect, only atypical *A. salmonicida* have been detected to date (Godoy et al., 2010).

Chile has been the second largest salmonid producer worldwide for several years, and salmon production is the second most important national economic activity (Carrasco-Bahamonde and Casellas, 2024). One of the leading causes for the Chilean salmon industry’s competitive disadvantage is the presence of infectious diseases. Piscirickettsiosis, a septicemic disease caused by the Gram-negative pathogen *Piscirickettsia salmonis*, is by far the most recurrent sanitary problem. Notwithstanding, the incidence of furunculosis in Atlantic salmon (*Salmo salar*) has shown a steady increase in recent years (Sernapesca, 2023). Although *A. salmonicida* had been detected in marine and fresh water cases several years ago in Chile (Godoy et al, 2010), freshwater disease has turned more prevalent since 2022, especially in aquaculture settings using recirculation technology. Therefore, the current epidemiological scenario supports the notion of re-emergence of furunculosis, as has already been suggested by some authors (Godoy et al., 2023). This situation can be ascribed to multiple causes such as production practices, host susceptibility, and control measures including antibiotic treatment and vaccination, but also to pathogen adaptation and evolution (Boerlin, 2022).

Several research groups have described molecular markers and developed PCR assays for the diagnostic of furunculosis (Gustafson et al., 1992; Byers et al., 2002; Beaz-Hidalgo et al., 2008a; Beaz-Hidalgo et al., 2008b; Rattanachaikunsopon and Phumkhachorn, 2012; Keeling et al., 2013; Fernandez-Alvarez et al., 2016; Bartkova et al., 2017a). One of the most widely used assays is based on the presence of *vapA*, a gene encoding the A-layer protein. This protein, which is also known as S layer, is a prominent virulence factor and immunodominant antigen of *A. salmonicida* species (Gustafson et al., 1992; Garduno et al., 1994; Lund et al., 2003). However, some doubts on the reliability of this PCR assay have been raised since the *locus* is prone to mutations (Gustafson et al., 1994; Lund and Mikkelsen, 2004; Vasquez et al., 2022). PCR protocols combining two or more markers can improve not only the detection capacity, but also broaden the scope of investigation to relevant genotypes associated with virulence and antibiotic resistance or antigenic patterns. In this regard, the development of a PCR protocol targeting the *fstA* gene, which codes for the ferric-siderophore receptor, has shown to be particularly effective and specific for the detection of *A. salmonicida* strains, with a negligible cross-reaction with related species (Beaz-Hidalgo et al., 2008b; Beaz-Hidalgo et al., 2013; Chapela et al., 2018). Of note, PCR protocols based on the detection of the *vapA* locus would not be expected to yield positive results in *A. salmonicida* subsp. *pectinolytica*, since it is lacking both *vapA* and the entirety of genes encoding the VapA protein secretion system necessary to assemble the A-layer (Merino et al., 2015)

Routine testing of some samples of presumed furunculosis cases that were delivered to our laboratory in 2022 resulted negative in *vapA-*based PCR assays. Taking into account the documented failures of this diagnostic tool, we hypothesized that the inclusion of a second target could help to improve the diagnostic performance. In line with the former, this work aims at determining the suitability of a *vapA-fstA* PCR scheme for the detection of *A. salmonicida*. We also applied the new assay to clinical tissue samples and detected *vapA-*absent bacterial types, which were isolated. Subsequent phenotypic characterization provided insights on culture features, *in vivo* virulence and antigenic profiles for this group of *A. salmonicida,* which nowadays represents a major proportion of furunculosis cases in Chile. Since *vapA* encodes a highly immunogenic protein, we also investigated if the current vaccination strategy is able to induce antibodies against the new strain, for which we examined the serological response in samples derived from fish under productive conditions.

## 2. MATERIALS AND METHODS

### 2.1. Clinical samples, bacterial strains and culture conditions

Tissues collected from suspected furunculosis cases were obtained from diagnostic routine, ADL laboratory, Puerto Montt, Chile. Samples, kept at -20 °C in preserving solution, were thawed to run a commercial RNA purification protocol (E.Z.N.A. Total RNA kit, Omega Bio-tek, GA, USA). Bacterial isolates used in this study were from the ADL strain collection, kept in 20 % glycerin (v/v) at -80 °C, and cultured on trypticase soy agar (TSA; BD, MD, USA) plates at 18 °C for 48 h. Some isolates were cultured in trypticase soy broth (TSB, BD). Dilutions prepared with saline were placed on TSA plates with Congo red added at 30 µg/ml before autoclaving the medium (Ishiguro et al., 1985). For subsequent molecular analysis, a loop of each isolate derived from a single colony was processed for DNA purification employing a GeneJET genomic DNA purification kit (Thermo Fisher Scientific, OR, USA). Epidemiological data of bacterial isolates are shown in Table 1.

**Table 1.** Epidemiological data related to bacterial strains. PCR results for *vapA* and *fstA* markers are shown as positive (p) or negative (n). Ss, *Salmo salar*; Om, *Oncorynchus mykiss*.

| # | Isolation date | Stage | Host | Company | Farm | Region | Water | Tissue | *vapA* | *fstA* |
| --- | --- | --- | --- | --- | --- | --- | --- | --- | --- | --- |
| AY-2485 | 02-13-13 | Adult | Ss | Company_1 | Farm_1 | XI | SW | Kidney | p | p |
| 2 | 03-14-13 | Adult | Ss | Company_1 | Farm_1 | XI | SW | Kidney | p | p |
| 3 | 03-14-13 | Adult | Ss | Company_1 | Farm_1 | XI | SW | Heart | p | p |
| 4 | 04-02-13 | Adult | Ss | Company_1 | Farm_1 | XI | SW | Spleen | p | p |
| 5 | 04-02-13 | Adult | Ss | Company_1 | Farm_1 | XI | SW | Heart | p | p |
| 6 | 01-10-14 | Adult | Ss | Company_1 | Farm_1 | XI | SW | Kidney | p | p |
| 7 | 01-10-14 | Adult | Ss | Company_1 | Farm_2 | XI | SW | Kidney | p | p |
| 8 | 01-23-14 | Adult | Ss | Company_2 | Farm_3 | XI | SW | Spleen | p | p |
| 9 | 03-24-16 | Fry | Ss | Company_3 | Farm_4 | XIV | FW | Skin lesion | p | p |
| 10 | 02-09-22 | Smolt | Ss | Company_4 | Farm_5 | X | FW | Kidney | p | p |
| 11 | 02-09-22 | Smolt | Ss | Company_4 | Farm_5 | X | FW | Kidney | p | p |
| 12 | 02-15-22 | Smolt | Ss | Company_4 | Farm_5 | X | FW | Skin lesion | p | p |
| 13 | 02-15-22 | Smolt | Ss | Company_4 | Farm_5 | X | FW | Kidney | p | p |
| 14 | 03-07-22 | Presmolt | Ss | Company_6 | Farm_5 | X | FW | Kidney | p | p |
| 15 | 03-09-22 | Juvenil | Ss | Company_7 | Farm_5 | X | FW | Kidney | p | p |
| 16 | 03-09-22 | Juvenil | Ss | Company_7 | Farm_5 | X | FW | Kidney | p | p |
| 17 | 03-09-22 | Juvenil | Ss | Company_7 | Farm_5 | X | FW | Skin lesion | p | p |
| 18 | 03-11-22 | Presmolt | Ss | Company_6 | Farm_5 | X | FW | Kidney | p | p |
| 19 | 03-11-22 | Presmolt | Ss | Company_6 | Farm_5 | X | FW | Kidney | p | p |
| 20 | 03-18-22 | Smolt | Ss | Company_4 | Farm_5 | X | FW | Kidney | p | p |
| 21 | 04-09-22 | Juvenil | Ss | Company_7 | Farm_5 | X | FW | Skin lesion | p | p |
| 22 | 04-09-22 | Juvenil | Ss | Company_7 | Farm_5 | X | FW | Gill | p | p |
| 23 | 04-25-22 | Smolt | Ss | Company_4 | Farm_5 | X | FW | Kidney | p | p |
| 24 | 04-28-22 | Smolt | Ss | Company_4 | Farm_5 | X | FW | Heart | p | p |
| 25 | 05-07-22 | Adult | Ss | Company_9 | Farm_7 | X | SW | Skin lesion | p | p |
| PM-118328 | 05-31-22 | Smolt | Ss | Company_6 | Farm_5 | X | FW | Kidney | p | p |
| 27 | 05-31-22 | Smolt | Ss | Company_6 | Farm_5 | X | FW | Kidney | p | p |
| PM-118565 | 06-20-22 | Fry | Ss | Company_6 | Farm_5 | X | FW | Skin lesion | n | p |
| 29 | 06-20-22 | Fry | Ss | Company_6 | Farm_5 | X | FW | Skin lesion | n | p |
| PM-119448 | 07-12-22 | Adult | Ss | Company_10 | Farm_8 | XII | SW | Kidney | n | p |
| 31 | 07-12-22 | Adult | Ss | Company_10 | Farm_8 | XII | SW | Kidney | n | p |
| 32 | 07-22-22 | Adult | Ss | Company_6 | Farm_9 | XI | SW | Spleen | p | p |
| 33 | 02-13-23 | Fry | Ss | Company_11 | Farm_10 | X | FW | Kidney | p | p |
| 34 | 02-13-23 | Smolt | Ss | Company_11 | Farm_10 | X | FW | Kidney | p | p |
| 35 | 02-17-23 | Fry | Ss | Company_11 | Farm_10 | X | FW | Kidney | p | p |
| 36 | 02-17-23 | Fry | Ss | Company_11 | Farm_10 | X | FW | Kidney | p | p |
| 37 | 02-18-23 | Fry | Om | Company_5 | Farm_11 | XIV | FW | Gill | p | p |
| 38 | 02-24-23 | Fry | Ss | Company_11 | Farm_10 | X | FW | Kidney | p | p |
| 39 | 05-18-23 | Adult | Ss | Company_4 | Farm_12 | XI | SW | Heart | p | p |
| 40 | 09-07-23 | Adult | Ss | Company_6 | Farm_13 | XI | SW | Kidney | p | p |
| 41 | 09-07-23 | Adult | Ss | Company_6 | Farm_13 | XI | SW | Kidney | p | p |
| 42 | 01-22-24 | Adult | Ss | Company_10 | Farm_14 | XII | SW | Kidney | n | p |
| 43 | 01-22-24 | Adult | Ss | Company_10 | Farm_14 | XII | SW | Kidney | n | p |
| 44 | 02-22-24 | Smolt | Ss | Company_11 | Farm_10 | X | FW | Skin lesion | n | p |
| 45 | 02-22-24 | Fry | Ss | Company_4 | Farm_5 | X | FW | Kidney | p | p |
| 46 | 02-22-24 | Fry | Ss | Company_4 | Farm_5 | X | FW | Kidney | p | p |
| 47 | 05-09-24 | Fry | Ss | Company_8 | Farm_6 | IX | FW | Skin lesion | n | p |

### 2.2 Bacterial growth kinetics, susceptibility to serum and counts

Selected isolates, representative of the different results obtained in *vapA* PCR assays (Table 1), were grown on TSA plates at 18 °C for 48 h. Then, a loop was used to prepare bacterial suspensions adjusted to optical density (OD_600_) = 1.00. Subsequently, 96-well microplates were inoculated with the bacterial suspensions diluted in TSB at 1:100. Bacterial growth kinetics of four *A. salmonicida* isolates were observed in an automatic spectrophotometer EPOCH (Biotek, VT, USA) at incubation temperatures of 10, 23, and 37 °C. Reads were taken up to 48 h. To determine the optimal salinity for growth, TSB was supplemented with 0, 0.5, and 2 % of NaCl. Aliquots of bacterial suspensions were log10 serially diluted in saline, plated in triplicates onto TSA, incubated at 18 °C for 48 h, and counted. For serum killing assays, a saline suspension of freshly cultured bacteria was prepared and adjusted to OD_600_ = 1.00. Samples were exposed to naïve serum collected from juvenile *S. salar* for 3 h at 18 °C. Bacterial counts were performed on control and treatment samples to compare the bactericidal action of the complement.

### 2.3 PCR assays and whole genome sequencing

*A. salmonicida* was identified in tissue samples and isolates using specific TaqMan qPCR assays targeting well-known virulence factors (Gulla et al., 2016a; Chapela et al., 2018). Samples were not treated with DNAse, hence, a combination of cDNA and gDNA was used as template. The sequences of primers and probes are listed in Table S1. PCR conditions were the following: 300 nM of each primer along with 200 nM of probe for each primer/probe set were mixed with 3 µl of nucleic acid template, enzyme and master mix using AgPath-ID™ One-Step RT-PCR and nuclease-free water according to the manufacturer’s instructions (Applied Biosystems™, TX, USA). To proceed with the amplification, 10 min at 45 °C for reverse transcription and 10 min at 95 °C for reverse transcriptase inactivation, then 45 cycles of 5 s at 95 °C and 30 s at 60 °C for annealing-extension were considered.

Whole genome sequencing was conducted at Codebreaker Bioscience facilities (Santiago, Chile). We first created a set of libraries using the standard protocol of Nextera Flex (Illumina™, CA, USA). Each standard Flex library was constructed using all standard kit reagents from the Nextera DNA Flex library prep kit, following the manufacturer’s protocol. The concentration of eluted libraries and the library size were measured using a Qubit high-sensitivity (HS) dsDNA kit (Thermo Fisher Scientific) and the S2 Cartridge from BIOtic with the Qsep1 Machine. Sequencing was performed on an Illumina™ Nextseq1000™ platform using a 600-cycle kit configured for 2×300 bp paired-end reads, incorporating a 5% PhiX mix as a diversity control.

### 2.4 Bioinformatic analysis

Raw fastq files were processed using a custom-developed Nextflow pipeline (https://github.com/gene2dis/mgap), which included quality control and cleaning of the raw reads with FastP (v.0.23.2) (Chen, 2023), followed by assembly using SPAdes (v.3.15.5) (Bankevich et al., 2012). We employed Bakta (v.1.7.0) for contig annotation (Schwengers et al., 2021) and BLASTN v2.15.0 to look for homologous sequences of contigs of interest in public databases (Altschul et al., 1990). For the visualization of groups of homologous biosynthesis gene clusters, the clinker tool was applied (Gilchrist and Chooi, 2021). Average nucleotide identities (ANI) were calculated using the ANI calculator tool hosted at https://www.ezbiocloud.net/ (Yoon et al., 2017).

A phylogenomic tree was generated by downloading all available *A. salmonicida* genomes that have been isolated from fish (as of January 2025), and filtering by genome completeness (>95 %) and contamination (<5 %) using CheckM2 (Chklovski et al., 2023). Filtered genomes were annotated with Bakta (Schwengers et al., 2021), and a pangenome analysis was performed using Roary (Page et al., 2015). The core genome, defined as genes shared by more than 95 % of all genomes, was aligned in MAFFTT (Katoh and Standley, 2013), and a phylogenetic tree was constructed with IQtree2 (Minh et al., 2020).

### 2.5 Purification and analysis of bacterial membrane antigens

A modified, Sarkosyl-based extraction strategy was applied to obtain the outer membrane protein (OMP) fraction (Maiti et al., 2011). Briefly, a loopful of isolates cultured on TSA was suspended in saline and centrifuged at 10,000 g for 10 min. The pellet was suspended in lysis buffer (300 mM NaCl, 10 mM HEPES, 2 mM PMSF, 8 M urea, pH 8.0), then sonicated on ice for 1 min at 70 % potency (300 V/T, Biologics Inc, VA, USA). The crude extract was centrifuged at 10,000 g for 10 min at 4 °C, resuspended in 1 ml of 0.2 % Sarkosyl and agitated over night at room temperature (RT). Samples were centrifuged at 10,000 g for 1 h at 4 °C, followed by pellet washing twice with 10 mM Tris-HCl pH 7.4 buffer and recovering the pellet by centrifugation at 10,000 g for 10 min at 4 °C. Finally, the OMP fraction was suspended in buffer containing 10 mM Tris-HCl pH 7.4 plus 8 M urea. Proteins were kept at -20 °C before use.

For LPS extraction and purification, we adapted a method described by others (Yi and Hackett, 2000). A loop of bacteria cultivated on TSA was suspended in 300 µl of Trizol reagent (Thermo Fisher Scientific) and incubated for 15 min at RT. Then, 100 µl of chloroform were added mixing thoroughly, followed by another incubation period of 10 min at RT. The water phase was centrifuged at 12,000 g for 10 min, then treated with 500 µl of 0.375 M MgCl_2_ in cold ethanol (-20 °C). The pellet was finally recovered by centrifugation at 12,000 g for 15 min at 4 °C. Membrane antigens were analyzed with discontinuous SDS-PAGE electrophoresis on 5 % and 15 % polyacrylamide stacking/solving gels. Gels were stained with Coomassie blue for protein analysis and silver-stained for LPS assessment according to a protocol previously described (Hitchcock and Brown, 1983).

For Western blots, a semi-dry protocol was followed (Bio-Rad, CA, USA). Signals in the blots were revealed using a polyclonal antibody prepared from serum of a rabbit immunized with a VapA*+* isolate (Aquit, https://aquit.net/) and the corresponding HRP-conjugated secondary antibody against salmon Ig heavy chain (LSBio, https://www.lsbio.com/).

### 2.6 Virulence *in vivo* testing

Juvenile Atlantic salmon, *Salmo salar* (mean weight 28 g) were intraperitoneally challenged with either one of three infectious doses of representative *A. salmonicida vapA*-absent PM-118565 field isolate (∼10^6^, ∼10^7^, or ∼10^8^ colony forming units [cfu]/fish) in independent tanks. Two tanks containing 500 L of freshwater, filtered and UV treated with a turnover rate of 1 h, were used for the assay. Each tank contained 120 fish in total: 30 healthy control fish (25 %) that received an intraperitoneal injection (ip) of 0.1 ml of saline, and 90 fish (75 %) which were injected with bacterial inoculum. This group of 90 fish was divided into three groups of 30 fish, as marked by Visible Implant Elastomer (VIE tagging), for one to receive 10^8^ cfu/fish and the other ones to be challenged with 10^7^ and 10^6^ cfu/fish, respectively. Biomass was adjusted to a density of ∼15 kg/m^3^ in one tank. In the second tank, we simulated overcrowding as a stress condition using 2× biomass density by diminishing the water level.

In parallel, we performed a lethal dose 50 (LD50) assay in a third tank containing 100 fish. These fish were inoculated with the *vapA*+ PM-118328 isolate at doses ranging from 10^4^-10^7^ cfu/fish, with each group comprising 20 animals. Accordingly, we distinguished five groups in this tank: the negative control group and fish that received 10^4^, 10^5^, 10^6^, and 10^7^ cfu, respectively. All experiments were carried out at 10.5±0.1 °C and terminated 30 days post-inoculation (dpi). Mortality was monitored and registered daily. Anterior kidney, liver and spleen pooled samples from each dead fish were obtained, placed in tubes with RNAlater (Thermo Fisher Scientific), and stored at -80 °C until further analysis. At 30 dpi, all survivor fish, including control fish, were sacrificed to take internal organ samples for PCR analysis. The experimental design was approved by ADL’s Bioethics Committee and performed in TEKBios fish trial center (study code 000-017).

### 2.7 Serum immune response

Immune responses to vaccination against *A. salmonicida* were evaluated in serum field samples. During vaccination activities that took place in farms 5, 7 and 10 in winter 2023 (Table 1), five unvaccinated *S. salar* pre-smolts were euthanized with an overdose of benzocaine, before being bled to obtain serum samples. Further serum samples were obtained from identical numbers of fish per farm during the freshwater stage at ∼300 and ∼600 thermal units (TU) time points post vaccination with a pentavalent product, including *A. salmonicida, P. salmonis, Vibrio anguillarum,* Infectious Pancreatic Necrosis virus (IPNV), and Infectious Salmon Anemia virus (ISAV) antigens. All serum samples were kept at -80 °C until processing. Briefly, 96-well microplates were coated using 5 µg/well of bacterial proteins purified from representative *A. salmonicida vapA+* or *vapA*-absent isolates (PM-118328 and PM-118565, respectively; see Table 1). Enzyme-linked immunosorbent assays (ELISA) were performed on protein-coated microplates containing 5 % skim milk as blocking agent, washed with phosphate buffer saline (PBS)-0.05 % Tween (PBS-T). Fish serum was added 1:100 in skim milk, followed by incubation for 1 h at RT. Microplates were subsequently washed with PBS-T, then incubated with a monoclonal HRP-conjugated antibody anti-salmon IgM, and washed again. Finally, the signal was revealed with the chromogenic substrate 3,3′,5,5′-tetramethylbenzidine (TMB; Thermo Fisher Scientific). Absorbance was captured at 450 nm with a spectrophotometer.

### 2.8 Statistical analysis and plots

Plots were constructed using the ggplot package on R (version 4.0.5). Differences in serum killing assays were assessed by t-test. For the *in vivo* assay, the percentages of cumulative mortalities were analyzed using the Kaplan-Meier method, and the differences were evaluated using the log-rank. For LD50 calculations, we used the BioRssay R package. Statistical tests were also performed with R.

## 3. RESULTS

### 3.1 Diagnostic statistics, tissue sample analyses and disease characterization

ADL Diagnostic’s routine database was interrogated for *A. salmonicida* testing and detection between January 2021 and June 2024. As shown in Figure 1A, a dramatic increase in the demand for *A. salmonicida* PCR diagnostics took place during 2022 in comparison with 2021. Since tissue samples from 2023 and earlier were analyzed in a single-marker qPCR assay, all positive cases were considered as caused by the *A. salmonicida vapA+* strain. During 2024, however, after the modification of the diagnostic routine and the introduction of the two-marker strategy, several cases of furunculosis tested positive only for *fstA* but not for *vapA*. Data revealed *A. salmonicida vapA-*absent types to account for 88 % of all tissue samples tested this year (Figure 1B). Typical clinical signs observed in field cases of *vapA-*absent furunculosis included skin lesions on the back, abdomen and fins, which were often coinfected with *Flavobacterium psychrophilum*. Signs of inflammation of internal organs, as usually described in *vapA*+ furunculosis, were rare in cases caused by *vapA*-absent types, but *vapA-*absent bacterial colonies were isolated from those samples (Figure 1C, Table 1).

**Figure 1.**
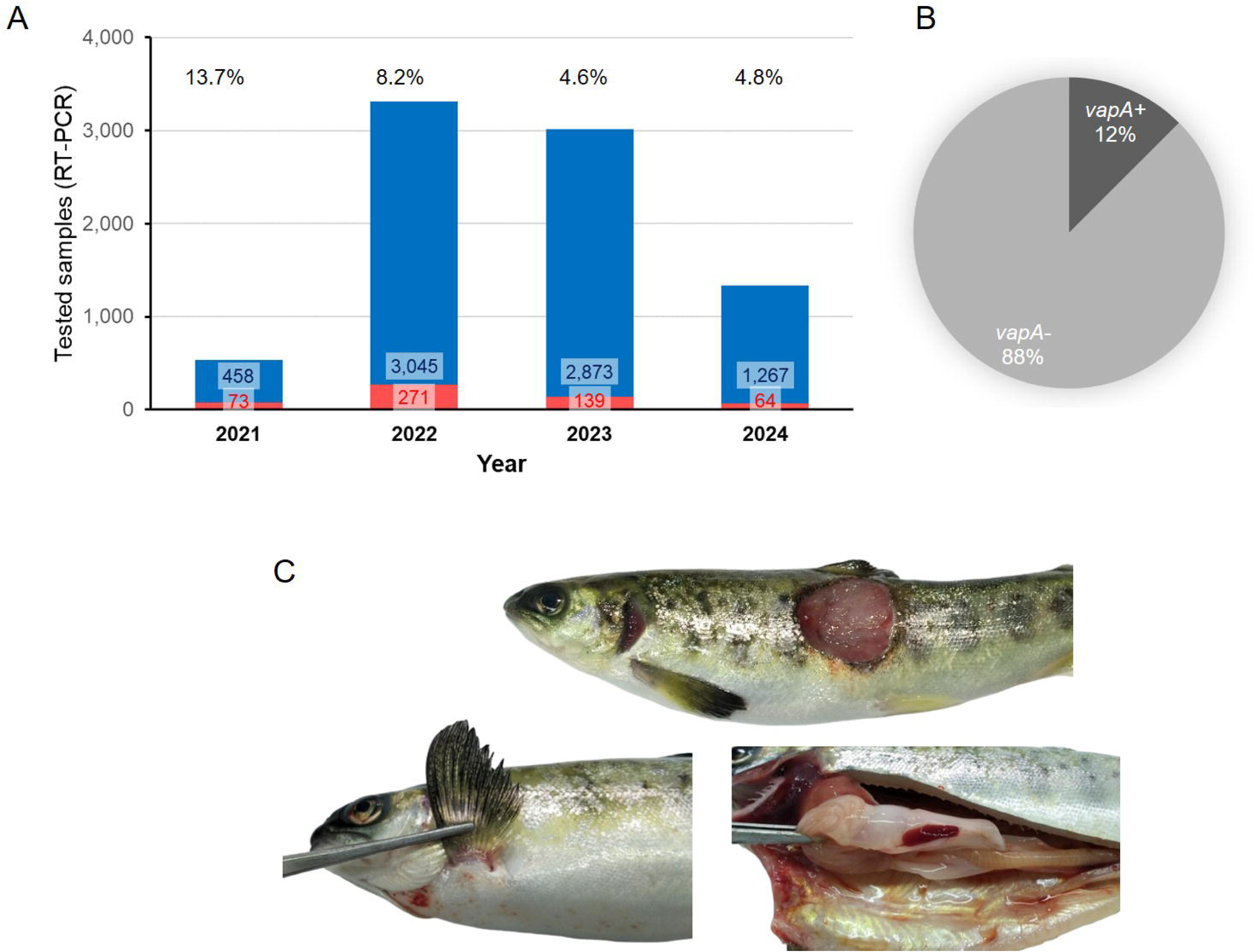
Statistics on past years’ *A. salmonicida* diagnostics in tissue samples (combination of cDNA and gDNA as template for PCR). A) Bar plot, number of positive samples and positive percentages are indicated per year. B) Cake plot, proportion of *A. salmonicida* isolates detected during 2024 (up to June, 30^th^) regarding their *vapA* genotype. C) Photographs of a representative furunculosis case caused by a *vapA-*isolate (ADL pathology records).

### 3.2 Bacterial isolation, genotyping and phenotypic profiling

ADL’s strain collection encompasses atypical *A. salmonicida* isolates that have been collected since 2013. Those old isolates stem from seawater cases, but the newer additions originate from freshwater outbreaks (Table 1). Bacterial specimens were isolated from diseased fish belonging to 11 different salmon farming companies located across five administrative regions. While they have their own farms, some companies utilize third-party facilities for rearing fish up to smolt size (*e.g*., Farm 5). Although some isolates did not amplify in *vapA* qPCR, they had been stored for further investigation due to their appearance and colony morphology. All isolates were genotyped for the presence of *vapA* and *fstA*. As shown in Table 1, the majority were *vapA*+/*fstA*+, with the *vapA*-/*fstA*+ genotype accounting for only 17% of the total number of isolates, all of which date to 2022 and later. This share is in disagreement with the prevalence detected in tissue samples, presumably because the number of processed samples and the success rate of isolation are not comparable to the PCR detection.

The A-layer phenotype was investigated using a classic culture method, taking advantage of the retention capacity of *vapA*+ colonies for Congo red (Ishiguro et al., 1985). As illustrated in Figure 2A, *A. salmonicida vapA*+ (represented by isolates AY-2485 and PM-118328) can adsorb the dye, forming deep red colonies, while those typed as *vapA*-/*fstA*+ were unable to retain the dye and remained gray (isolates PM-118565 and PM-119448). Since the retention of Congo red depends on the expression of the A-layer protein, these observations allow us to infer that *vapA*-absent colonies are devoid of this molecule. Another important phenotypic feature we assessed was the growth at different temperatures and salinities. Growth curves obtained demonstrate optimal growth for all isolates at mesophilic temperatures (23 °C) at different salinity levels, except for *vapA+* AY-2485 at high salinity. All *A. salmonicida* isolates showed the capacity to adapt to psychrophilic temperatures (10 °C) and mild salinity, with the exception of AY-2485, which did not replicate. Attempts to define conditions that enable better growth of AY-2485 failed, and this isolate behaved persistently as a slow-growing, fastidious bacterium (Figure S1). Growth at 37 °C, albeit suboptimal, could be observed for *vapA-*absent genotype isolates only. In this context, high salinity proved to be a detrimental factor for growth for PM-118565 but not PM-119448 (Figure 2B). Regarding these isolates’ susceptibility to naïve fish serum, *vapA*-absent isolates – which are devoid of the A-layer – resulted more susceptible to being killed than those carrying an intact copy of *vapA* (Figure 2C).

**Figure 2.**
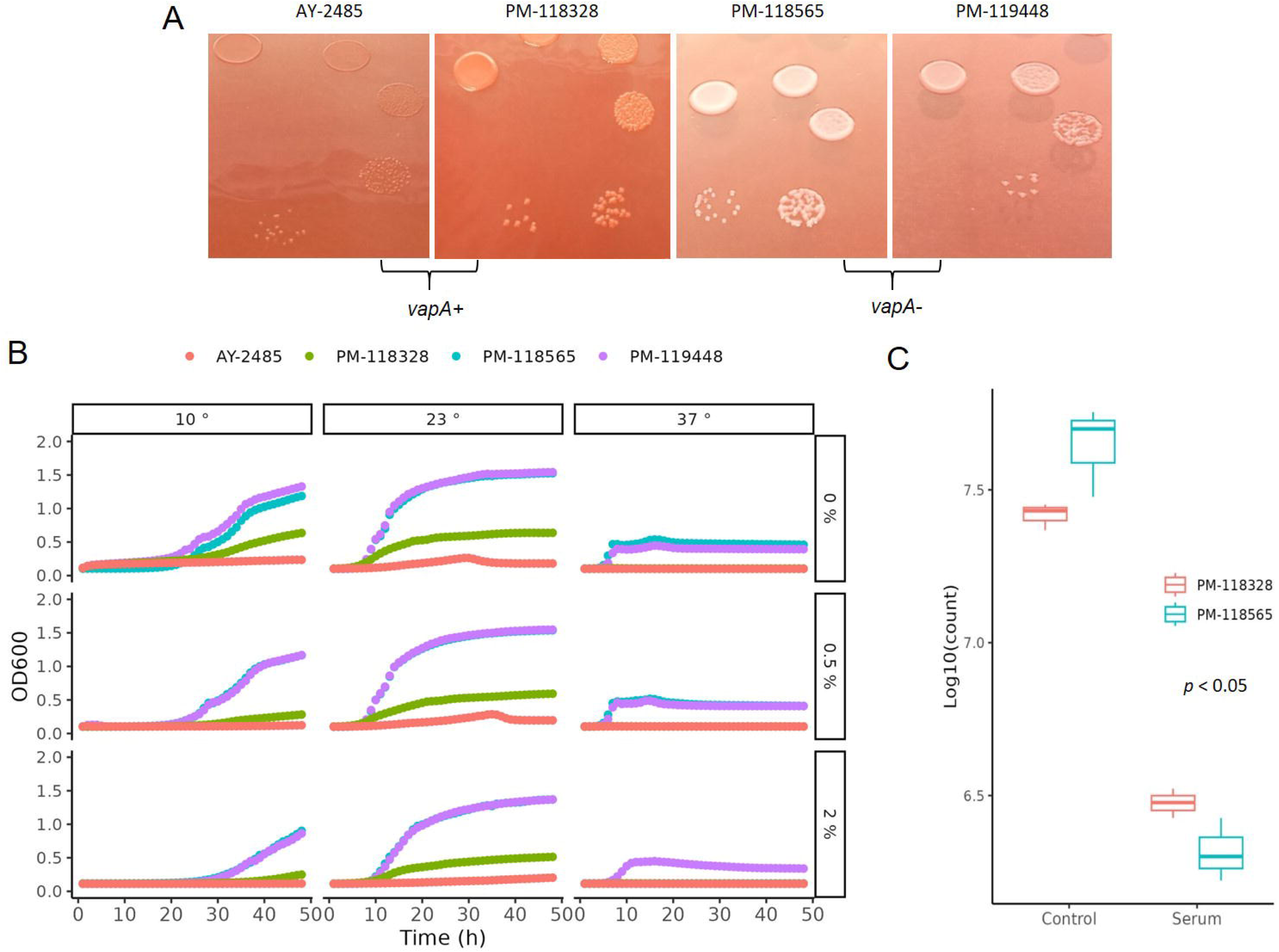
Phenotypic characterization of *A. salmonicida* VapA-isolates. A) Bacterial colonies on TSA plus Congo red. B) Growth kinetics on TSB at different temperatures and salinities. C) Serum killing assays.

### 3.3 Phylogenetic and bioinformatic analysis

The sequences of all draft genomes were deposited in GenBank under the BioProject PRJNA1168199. The corresponding BioSample accession numbers along with data of draft genomes are listed in Table 2. Genome-based phylogeny placed sequences of *vapA+* PM-118328 and AY-2485 isolates within a cluster of *A. salmonicida* sequences derived from diseased fish, mostly from salmonids (Figure 3, Table S2). Interestingly, PM-118328 grouped together with sequences derived from reared salmon in Canada, while AY-2485 forms a single-unit subcluster. In contrast, *vapA*-absent types clustered adjacent to the sequences derived from non-salmonid fish and environmental samples, suggesting a different lineage compared to Chilean *vapA+* types.

**Figure 3.**
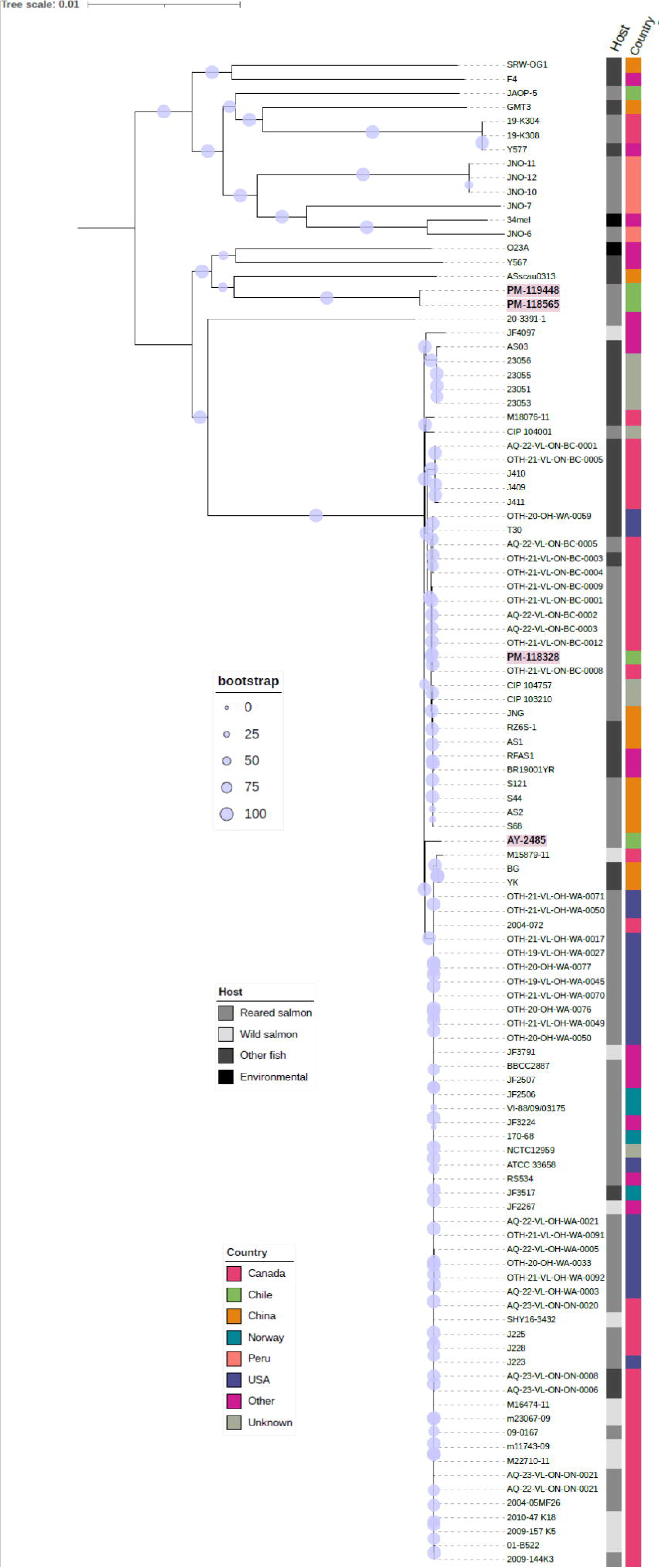
Phylogenomic analysis. Unrooted consensus tree based on the genome sequences of *A. salmonicida* isolates recovered from diseased fish (bootstrap level = 1,000 replicates). The percentage of replicate trees in which the associated taxa clustered together in the bootstrap test are shown next to the branches using a circle scale. Only values >50 % are shown. Sequenced isolates from this work are written in bold. Variables “Country” and “Host” are depicted using colored and grey labels, respectively.

**Table 2.** BioSample accessions and sequencing data for draft genomes.

| Accession | Isolate | Organism | Tax ID | Contigs | Length<br>(Mb) | GC (%) | *vapA* |
| --- | --- | --- | --- | --- | --- | --- | --- |
| SAMN44032016 | PM-118565 | *A. salmonicida* | 645 | 545 | 4.71 | 58.85 | - |
| SAMN44032017 | PM-119448 | *A. salmonicida* | 645 | 827 | 4.99 | 58.72 | - |
| SAMN44032018 | PM-118328 | *A. salmonicida* | 645 | 1949 | 5.02 | 57.99 | + |
| SAMN44032019 | AY-2485 | *A. salmonicida* | 645 | 493 | 4.61 | 58.50 | + |

A comparative analysis of draft genomes at a large scale is presented in Table 3. We performed a pair-wise genome comparison using an improved algorithm to calculate ANI. Even though *vapA+* isolates were collected with a decade of difference, their sequences exhibited the highest identity value (99.59 %). The identity between mixed group-pairs resulted to be over 97 %. On the contrary, the comparison of *vapA*-absent genomes yielded even higher identity scores, reaching 99.96 % of identity, despite them not being epidemiologically related. Comparative analysis performed on the *vapA* locus and its genomic context disclosed a striking result: As expected, we could not find homologous sequences for *vapA* in PM-118565 and PM-119448 genomes, which is consistent with the PCR results. Notably, further contig and annotation analysis allowed us to unveil a fragment encoding a capsule biosynthesis gene cluster flanked by some O-antigen biosynthesis *wb*_salmo_ genes in *vapA-*absent types (Figure 4). This region is quite different not only in *vapA+* types, in which the *locus* harbors genes of *wb*_salmo_, *vapA* and its protein-exporting machinery, but also in the *vapA-*absent SRW-OG1 genome. The genomic organization found in *vapA+* isolates resembles that described for well-known *A. salmonicida* strains (Merino et al., 2015). According to the synteny analysis, it seems that a complex sequence rearrangement comprising *rfb* genes took place in *vapA*-absent types, since that cluster appears to be split into *rfbBA* and *rfbDC*, which are flanking the capsule biosynthesis cluster instead of presenting as a continuous sequence of elements. Evidence on the presence of the same *locus* organization was collected by means of BLAST searches, which identified an 14-kb orthologous region in *Aeromonas* sp. O23A, an environmental strain related to *A. salmonicida* (Uhrynowski et al., 2017).

**Figure 4.**
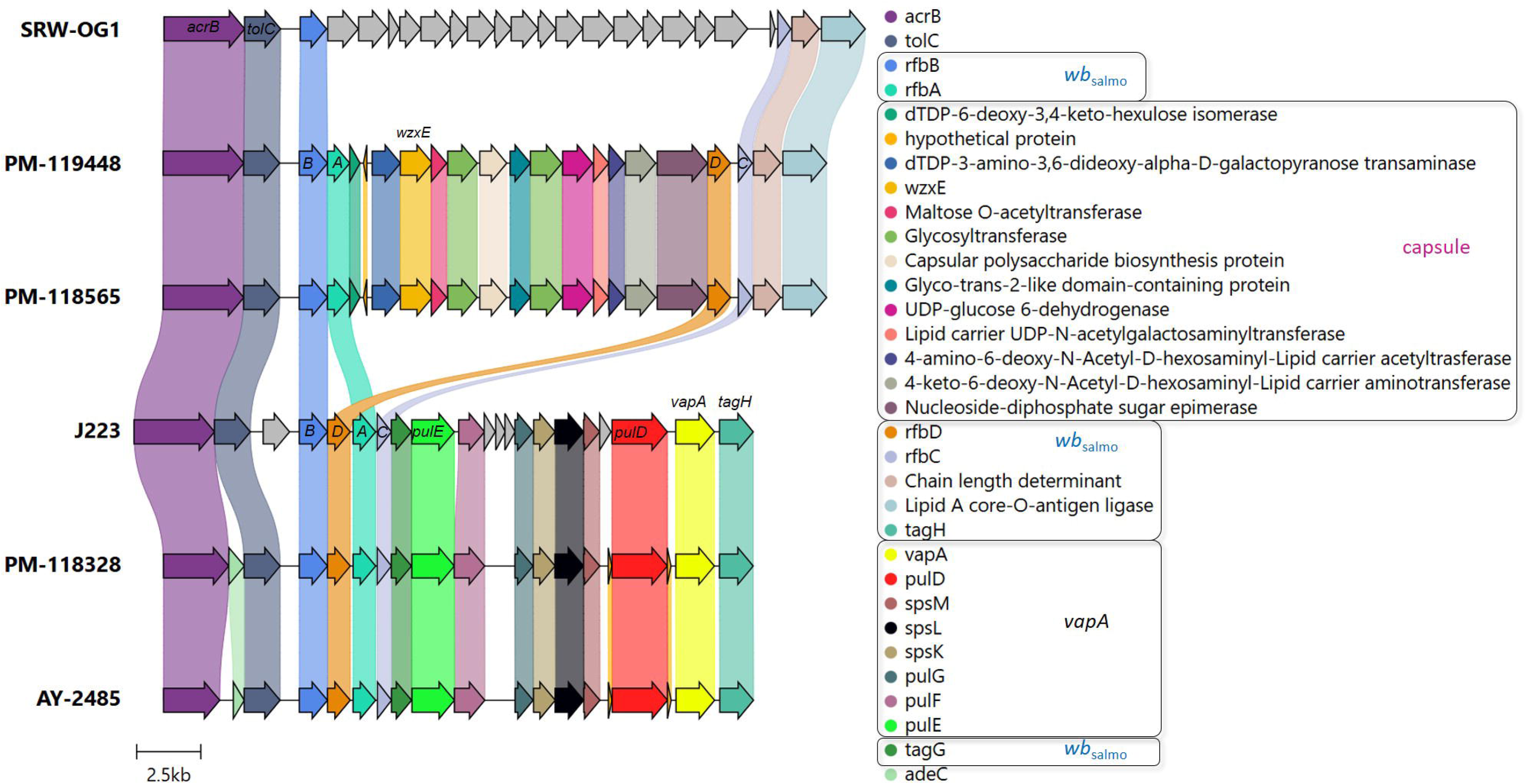
*vapA* locus organization in *vapA*-absent and *vapA+* isolates. Arrows depicting genes and bullet points preceding the gene names follow the same color code. Coding sequences related to the *wb*_salmo_ and *vapA* clusters (for details, see Merino et al., 2015) along with the capsule biosynthesis gene cluster predicted in *vapA*-absent isolates appear within black squares in the legend. Some gene names are depicted to aid interpretation. *ABCD* correspond to the *rfb* gene cluster. J223 and SRW-OG1 refer to *vapA*-related *A. salmonicida* phenotypes studied in the present work. Annotation details for the cluster found in the SRW-OG1 chromosome are shown in Table S4.

**Table 3.**
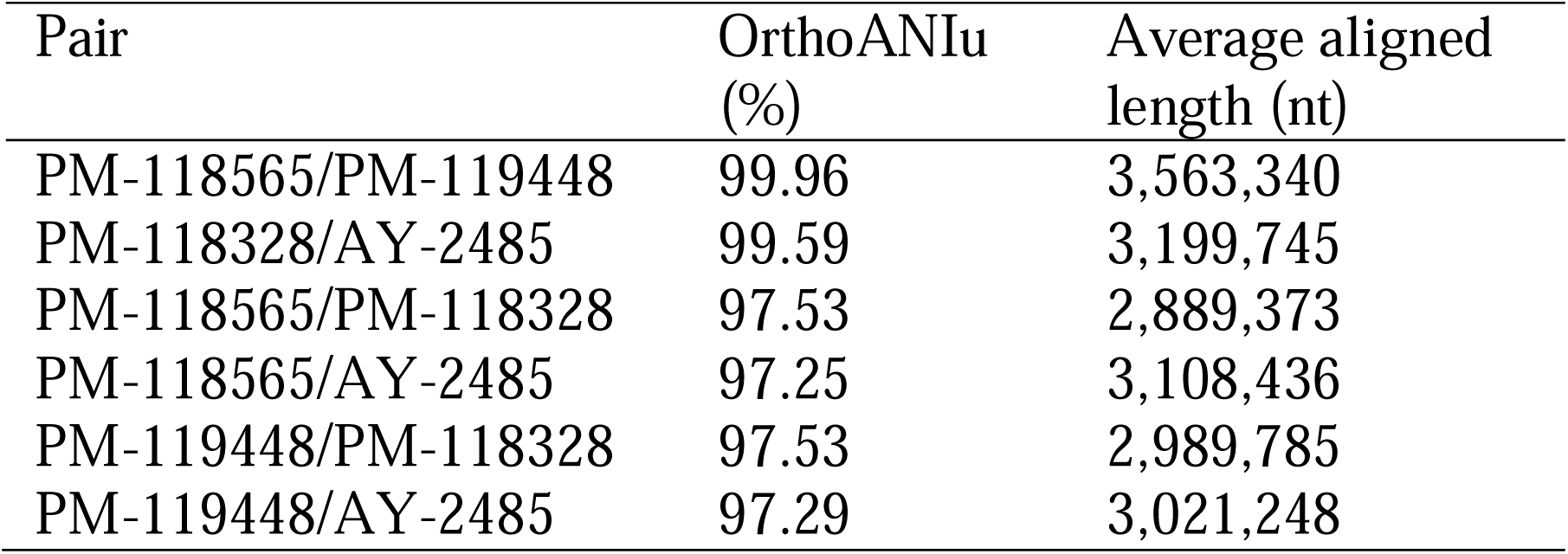
Results of ANI analysis.

### 3.4 Membrane antigen profiling

Outer membrane protein profiles disclosed specific patterns for both *vapA*-absent and *vapA+ A. salmonicida* isolates (Figure 5A). Further analysis concerning the protein fraction reactive to rabbit immunized serum revealed some common bands but most notably confirmed the absence of a ∼49-kDa band compatible with the VapA protein in *vapA*-absent isolates (Figure 5B). Isolates could also be grouped according to their LPS profiles: *vapA*-absent types presented a single-band pattern which is structurally consistent with the lipid A-core oligosaccharide. By contrast, *vapA*+ isolates exhibited two bands. One of them probably corresponds to the lipid A-core (low molecular size), while the other likely represents the lipid A-core plus O-antigen (high molecular size) (Figure 5C). LPS Western blots evidenced a high-molecular size single-band pattern in *vapA+* isolates (Figure 5D). This finding is in accordance with the above description of the LPS profiles.

**Figure 5.**
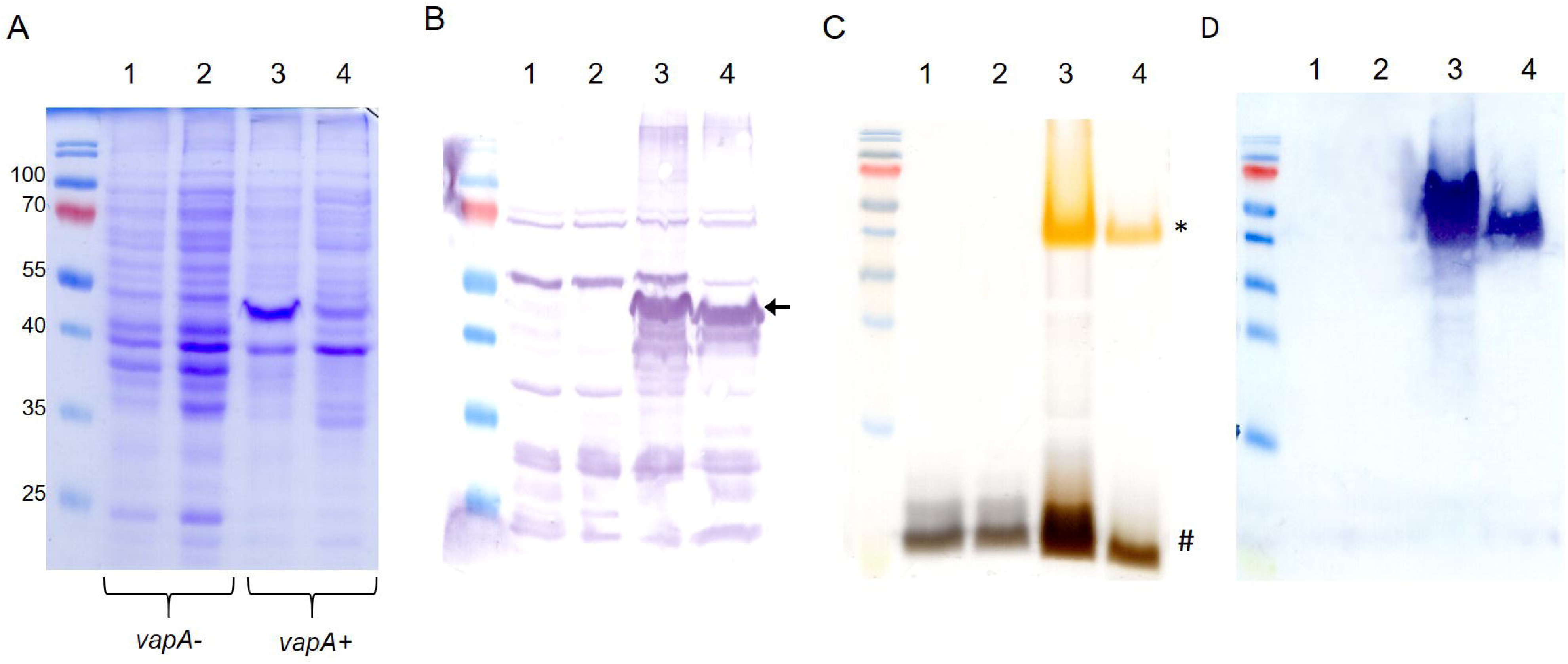
Outer membrane components profiling of *A. salmonicida* isolates. A) OMP profiles. B) OMP Western blot using serum from a rabbit hyperimmunized against VapA+ *A. salmonicida*. The black arrow indicates the band corresponding to the putative VapA protein. C) LPS patterns in silver-stained SDS-PAGE. Low and high LPS molecular species are indicated with # and *, respectively. D) LPS Western blot revealed with the same antibody used in B. Lanes 1-4 correspond to samples derived from PM-118565, PM-119448, PM-118328, and AY-2485 isolates, respectively.

### 3.5 Virulence

Remarkable differences in virulence between *vapA*-absent and *vapA+* isolates became evident in the intraperitoneal challenge model of infection. The *vapA*+ strain killed fish in a few days: Mortality rates of 100 % were reached as fast as five dpi in *S. salar* inoculated with the highest bacterial dose (3.17 × 10^7^ cfu/fish, Figure 6A). The same level of mortality was recorded at six dpi in fish challenged with the lower dose of 2.67 × 10^6^ cfu, while 90 and 65 % mortality were accumulated with 1.50 × 10^5^ and 1.80 × 10^4^ cfu/fish, respectively. In contrast, mortality observed in fish infected with the *vapA*-absent PM-118328 strain scaled up to 80 and 6.7 % after the inoculation of 4.0 × 10^8^ and 3.67 × 10^7^ cfu/fish, respectively (Figure 6B, left panel). Interestingly, the same bacterial doses induced 90 and 23 % mortality in fish that experienced overcrowding (Figure 6B, right panel). The relative bacterial loads in kidney, liver, and spleen pooled samples were determined by PCR for all fish, mortality and survivors. Results show that both *vapA*+ and *vapA-*absent isolates did infect internal organs, producing the death of fish most probably from a septicemic disease (Figure 6C). According to calculated LD50 values, virulence ranked as *vapA+* > *vapA-*absent+stress ≈ *vapA-*absent (Figure S2).

**Figure 6.**
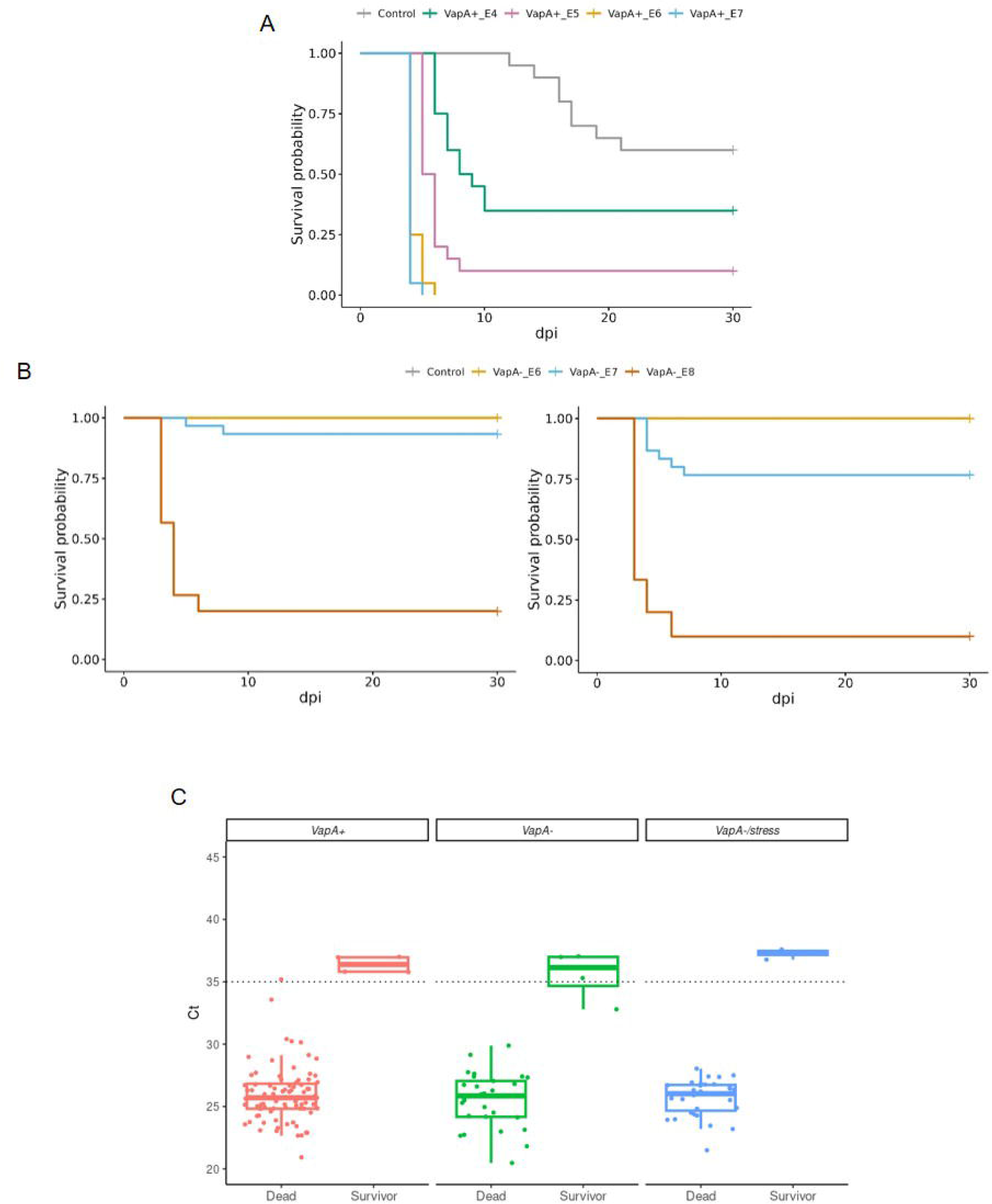
Virulence *in vivo* of *A. salmonicida* isolates in juvenile *S. salar*. A) Survival curves for fish inoculated with VapA+ PM-118328 at ∼10^7^ (VapA+_E7), ∼10^6^ (VapA+_E6), ∼10^5^ (VapA+_E5) or ∼10^4^ (VapA+_E4) cfu/fish, or saline (Control). B) Left panel, survival analysis of fish inoculated with VapA-PM-118565 using ∼10^8^ (VapA-_E8), ∼10^7^ (VapA-_E7), ∼10^6^ (VapA-_E6) cfu/fish, or saline (Control). Right panel, results for the same challenge doses in fish subjected to overcrowding as a stress condition. Some curves appear overlapped at survival probability= 1.00. C) Relative bacterial loads in internal organs measured by *fstA* PCR. Box and whisker plots represent all groups of fish tested, including controls. Dotted line indicate the threshold value (Ct=35).

### 3.6 Immune response of vaccinated fish

Relative IgM serum levels against *A. salmonicida* were determined by ELISA. As expected, vaccinated fish exhibited an increasing level of antibodies during the development of immunity in fresh water (up to ∼600 TU), which reflects the homologous character of the antigen present in the vaccine formulation (VapA+-coated microplates, Figure 7). A different situation was observed in farm 7, where fish presented a high level of anti-*A. salmonicida* IgM prior to the administration of the vaccine. These fish’ specific IgM serum levels dropped to basal values at ∼600 TU. When the same samples were tested on microplates coated with VapA-extract, very low antibody levels were detected, with the exception of fish from farm 10, which disclosed a behavior similar to that obtained with VapA+-coated microplates, albeit at a lower relative level (Figure 7).

**Figure 7.**
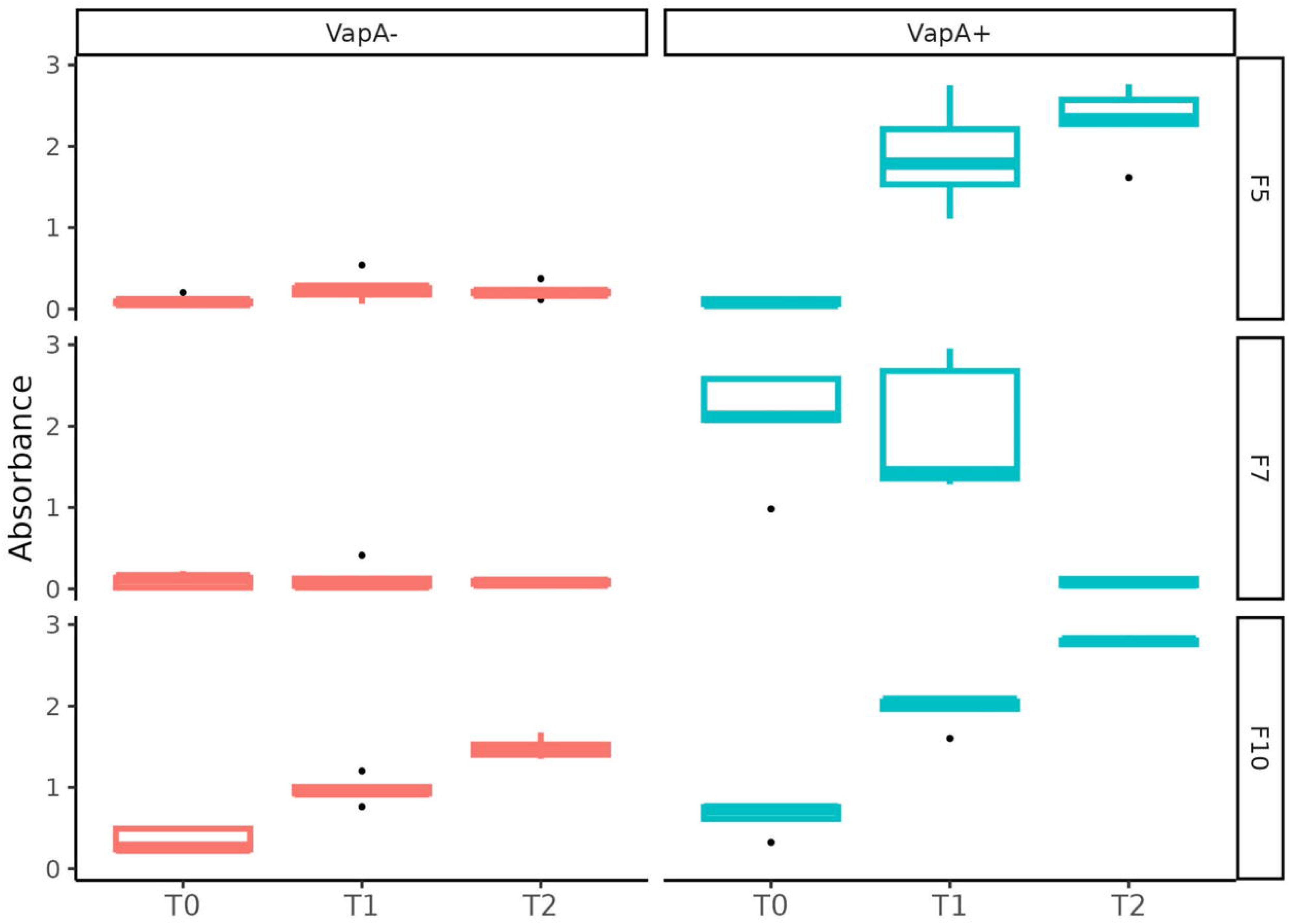
Serum immune response against *A. salmonicida* isolates determined by ELISA. Fish derived from three farms (F5, F7 and F10) were sampled at 0 (T0), ∼300 (T1), ∼600 (T2) TU after vaccination. Relative IgM levels against *A. salmonicida* were examined in independent microplates coated with VapA+ or VapA-protein extracts.

## 4. DISCUSSION

Some bacterial pathogens have developed a series of structures to avoid recognition by the host’s immune system and resist defense mechanisms. This also includes the virulence array protein gene *vapA*, which plays a crucial role in the pathogenicity and differentiation of various subspecies of *A. salmonicida* (Gulla et al., 2016b). This is the first report demonstrating that, contrary to what we would have expected, a strain lacking the A-layer is mainly responsible for recent furunculosis outbreaks in Chile. According to our results, a second PCR marker for the detection of *A. salmonicida* not only improved the sensitivity of our diagnostic scheme, but also allowed us to identify a neglected strain, adding a layer of complexity to the epidemiology of the pathogen. In this regard, we must note that we cannot currently assess the extent to which cutaneous co-infection with *F. pyschrophilum* may affect the course of atypical furunculosis, though the lack of dissemination to internal organs argues against a major pathophysiological role and rather suggests a link to environmental exposure. In a global context, furunculosis caused by *vapA*-absent *A. salmonicida* isolates is not unheard of, as recent reports of furunculosis caused by phenotypically similar isolates of *A. salmonicida* in juvenile Siberian sturgeons (*Acipenser baerii*) and in reared snakehead fish (*Channa argus*) show (Sun et al., 2023; Vazquez-Fernandez et al., 2023). On the other hand, since there are no tissue samples stored in our biobank derived from furunculosis cases before 2024, we cannot clarify when the *vapA*-absent strain arose, and it is also hard to predict if this shift in prevalence will prevail over time. Our evidence suggests that the *A. salmonicida vapA*-absent strain is turning dominant and has likely overcome its *vapA+* counterpart very recently. We should also keep in mind the possibility that additional *A. salmonicida* strains or species-related isolates could fail to be identified by the PCR scheme proposed in this work. In fact, our strain collection bears some *A. salmonicida*-like isolates whose colonies look similar to those analyzed in this study, but which do not amplify *vapA* or *fstA* (data not shown). Sequencing studies in progress will permit insights on the taxonomic affiliation of these isolates.

The fact that some *A. salmonicida* isolates do not produce the A-layer has been linked to the heterogeneity of the *vapA* locus (Lund and Mikkelsen, 2004; Gulla et al., 2016b), with *A. salmonicida* subsp. *pectinolytica* being the only subspecies that naturally occurs with this membrane phenotype. Due to its relevance in virulence and antigenicity, the absence of this gene could be related to adaptive changes resulting from host-pathogen interactions, thus favoring a relationship with certain advantages for the latter. Indeed, *A. salmonicida* subsp. *pectinolytica* is notable for lacking both the *wb*_salmo_ and A-layer (*vapA*) genes. The absence of these two major virulence factors in this atypical subspecies suggests a reduced virulence, making it less likely to cause disease in fish (Merino et al., 2015). Our results also highlight an important difference between *A. salmonicida* subsp. *pectinolytica* and Chilean *A. salmonicida vapA-*absent types, namely the replacement of the *wb*_salmo_ and *vapA* loci by a capsule biosynthesis gene cluster. This finding can be extended to all *vapA*-absent types analyzed in this work, as indicated by an additional PCR assay targeting the capsular polysaccharide biosynthesis protein, a gene located in this region (Figure 4, Table S3). Interestingly, the capsule biosynthesis gene cluster is also present in the genome of *Aeromonas* sp. O23A, an environmental strain isolated from arsenic-contaminated mine sediment in Poland, which seems to be well-adapted to living in such a harsh habitat, but may be transitioning from a former pathogen life style (Uhrynowski et al., 2017).

Genetic evidence is consistent with the LPS single-band pattern detected in the *A. salmonicida vapA*-absent strain. It probably corresponds to lipid A-core motifs and suggests that these isolates’ LPS molecule lacks the polymeric O-antigen (Figures 5C and 5D). Yet, it remains unclear whether the *vapA*-absent strain carries a functional capsule that contributes to their pathogenicity and antigen masking ability. The presence of O-antigen has been shown to be a reliable predictor of virulence: O-antigen-devoid isolates are less virulent in comparison with related strains that have complete LPS (Lerouge and Vanderleyden, 2002). In the present study, the behavior displayed by the *vapA-*absent strain during the *in vivo* testing is in accordance with a reduced virulence in comparison with *vapA+* types (Figure 6A). In fact, mortality was recorded at higher doses of the pathogen via ip injection, while the bacterium was unable to horizontally disseminate towards the cohabitant control group (Figure 6B). Taking into account what we know about the role of the A-layer and LPS in terms of *A. salmonicida* virulence, our results support the hypothesis that the virulence phenotype of *vapA-*absent types cannot solely be justified by a missing A-layer, but is also related to an incomplete LPS molecule. This finding makes sense in the light of the current epidemiological situation of furunculosis: outbreaks are often linked to specific processes in fish-rearing management, such as vaccination procedures or transfer to seawater, and result in limited mortality. Our results also suggest that *vapA*-absent types may induce higher mortality in fish subjected to suboptimal rearing conditions, although this finding did not reach statistical significance (Figure S2). Hence, it seems that *vapA-*absent types need stressful conditions to become a clinical issue.

The phenotypic characteristics exhibited by *vapA*-absent isolates are worth mentioning in more detail. According to the literature, the bacteria characterized in the present study are clearly mesophilic since they were primary isolated at 18 °C, grew well at 23 °C, showed potential to grow at psychrophilic temperatures, and even at 37 °C (Vincent and Charette, 2022). These features, along with these isolates’ capacity to adapt to a salinity gradient, are consistent with a versatile lifestyle (Gustafson et al., 1994; Chu et al., 1995). While the mechanism behind that adaptive capacity was beyond the scope of this study, the flexibility in the growth requirements of *vapA-*absent isolates argues in favor of their ubiquity, supporting the ability to transit between fresh and seawater salmon-rearing settings. The spread of *A. salmonicida* mesophilic strain with a degree of divergence from those atypical subspecies in Chilean aquaculture settings is contrasting with the current situation reported in other countries, in which recent, temporally unrelated furunculosis outbreaks were caused by *A. salmonicida* subsp. *salmonicida* isolates with over 99.9 % of identity, suggesting a single epidemiological unit (Wojnarowski et al., 2024). As mentioned above, our results support the current epidemiological scenario in Chile to be explained by at least two *A. salmonicida* strains. This conclusion may even extend to three strains, considering the particular growth requirements and phylogenomic placement of the AY-2485 isolate.

In Chile, the most popular vaccination strategy is based on a pentavalent product, which is administered in pre-smolt fish (Flores-Kossack et al., 2020). Current authorized vaccines comprising *A. salmonicida* antigen were developed more than 10 years ago, and it is plausible to infer that these formulations only include *A. salmonicida* VapA*+* strains. This can be deduced from the immune response determined by ELISA (Figure 6). Fish serum, regardless of its origin, showed a characteristic curve for adaptive immune response development when tested on microplates retaining the specific anti-VapA IgM. Since both strains share common OMP immunogenic profiles, with the exception of the VapA protein, we can speculate that IgM levels observed may be partially explained by the presence/absence of this antigen. This is supported by the literature, where VapA has been described as forming a layer array covering the outer membrane, being immunodominant and acting as protective antigen in vaccine formulations (Lund et al., 2003; Ebanks et al., 2005). Yet, the O-antigen is also recognized as immunodominant (Wang et al., 2005), and we cannot rule out the contribution of the O-antigen to the IgM levels. This assumption is supported by the fact that only a complete LPS, as extracted from *vapA*+ types, was able to capture specific antibodies. *A. salmonicida* LPS is indeed intriguing: LPS from Gram-negative pathogens, including well-known fish pathogens such as *P. salmonis*, triggers the production of antibodies (Herrera et al., 2022), however, it seems that *A. salmonicida*’s lipid A-core does not exhibit immunogenic properties, an observation also made by others (Wang et al., 2007). Further research is needed to elucidate if this may be part of a pathogen evasion mechanism.

## 5. REMARKS

Our results support the idea that a new *A. salmonicida* strain has emerged and become highly prevalent in Chilean salmon farms. Furthermore, its particular outer membrane antigen composition seems to be related to a mild course of disease, but may also contribute to reduced vaccine effectivity. Due to the relevance of furunculosis in sanitary programs, it should be stressed that a thorough functional characterization along with a molecular screening is required for an effective disease surveillance and control. Ongoing research will help to define the need for revising the vaccine strategy and implementing improvements.

## DATA AVAILABILITY STATEMENT

The original contributions presented in the study are included in the article/Supplementary Material. Data supporting the findings of this study are available on request from the corresponding author. Further inquiries can be directed to the corresponding author.

## ETHIC STATEMENT

The protocol for fish trials was reviewed and approved by ADL Diagnostic Chile bioethics committee.

## ACKNOWLEDGMENTS

This study was supported by the Chilean Economic Development Agency, CORFO, through grant PI-5297, and by internal funds from ADL Diagnostic Chile and TEKBios.

